## Supplementary data for "The encephalomyocarditis virus Leader promotes the release of virions inside extracellular vesicles via the induction of secretory autophagy"

**Supplementary figure 1**

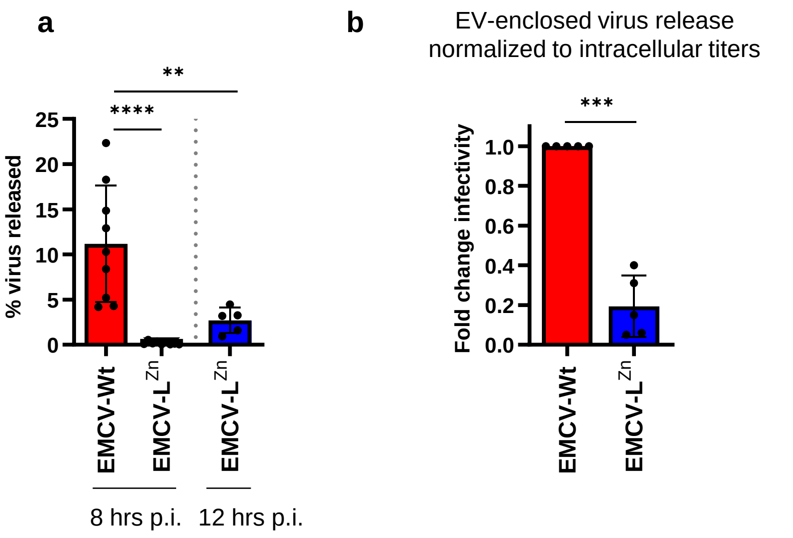

Supplementary figure 2

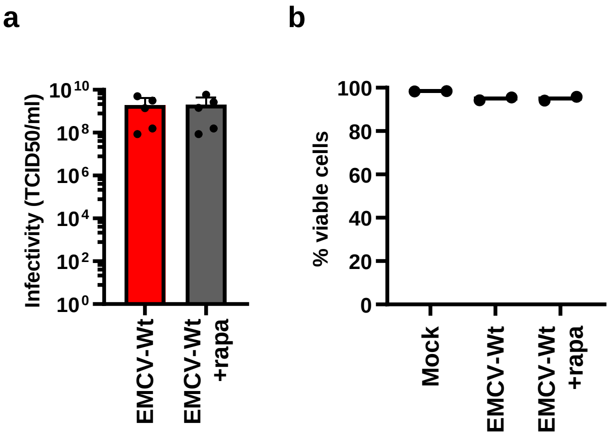

**Supplementary figure 3**

**a**

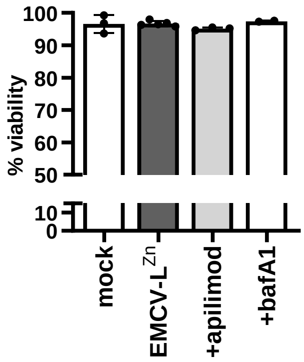

**Supplementary table 1**

**Proteins shared between EMCV-Wt & EMCV-L$^{Zn}$ & mock (n=181)**

| Accession | Gene | Accession | Gene | Accession | Gene | Accession | Gene |
|---|---|---|---|---|---|---|---|
| P60709 | ACTB | Q08554 | DSC1 | P14923 | JUP | P62826 | RAN |
| Q562R1 | ACTBL2 | Q02413 | DSG1 | P32004 | L1CAM | P62834 | RAP1A |
| P68032 | ACTC1 | P15924 | DSP | P11279 | LAMP1 | P61224 | RAP1B |
| P12814 | ACTN1 | P60981 | DSTN | P00338 | LDHA | P35241 | RDX |
| O43707 | ACTN4 | P68104 | EEF1A1 | P07195 | LDHB | P61586 | RHOA |
| P02768 | ALB | P13639 | EEF2 | P49257 | LMAN1 | P62913 | RPL11 |
| P04075 | ALDOA | P06733 | ENO1 | P02788 | LTF | P61353 | RPL27 |
| P09972 | ALDOC | P15311 | EZR | P61626 | LYZ | P62888 | RPL30 |
| P04083 | ANXA1 | P00742 | F10 | P20645 | M6PR | P18124 | RPL7 |
| P50995 | ANXA11 | Q5D862 | FLG2 | P29966 | MARCKS | P60866 | RPS20 |
| P07355 | ANXA2 | Q16658 | FSCN1 | P49006 | MARCKSL1 | P23396 | RPS3 |
| P08758 | ANXA5 | P06241 | FYN | P43121 | MCAM | P62070 | RRAS2 |
| P02656 | APOC3 | P04406 | GAPDH | P40925 | MDH1 | P31949 | S100A11 |
| P61204 | ARF3 | O14556 | GAPDHS | P14174 | MIF | P06703 | S100A6 |
| P18085 | ARF4 | P50395 | GDI2 | P26038 | MSN | Q8WTV0 | SCARB1 |
| P05089 | ARG1 | P29992 | GNA11 | P60660 | MYL6 | P01008 | SERPINC1 |
| P52565 | ARHGDIA | Q14344 | GNA13 | Q12965 | MYO1E | P05546 | SERPIND1 |
| P00966 | ASS1 | P04899 | GNAI2 | P15531 | NME1 | P31947 | SFN |
| P05023 | ATP1A1 | P08754 | GNAI3 | Q9Y639 | NPTN | P53985 | SLC16A1 |
| P13637 | ATP1A3 | P50148 | GNAQ | P01111 | NRAS | O15427 | SLC16A3 |
| P05026 | ATP1B1 | P63092 | GNAS | Q9UKS6 | PACSIN3 | Q15758 | SLC1A5 |
| P54709 | ATP1B3 | P62873 | GNB1 | Q9Y6V0 | PCLO | P11166 | SLC2A1 |
| P20020 | ATP2B1 | Q9HAV0 | GNB4 | O75340 | PDCD6 | P11169 | SLC2A3 |
| Q01814 | ATP2B2 | Q9UBI6 | GNG12 | Q8WUM4 | PDCD6IP | P08195 | SLC3A2 |
| P25311 | AZGP1 | P06744 | GPI | P30101 | PDIA3 | Q8NBI5 | SLC43A3 |
| P80723 | BASP1 | Q8NFJ5 | GPRC5A | P07737 | PFN1 | Q01650 | SLC7A5 |
| P35613 | BSG | P69905 | HBA1 | P18669 | PGAM1 | P27105 | STOM |
| Q10589 | BST2 | P68871 | HBB | P00558 | PGK1 | Q16563 | SYPL1 |
| P62158 | CALM1 | P01891 | HLA-A | O43175 | PHGDH | P02786 | TFRC |
| P31944 | CASP14 | P30493 | HLA-B | P14618 | PKM | P60174 | TPI1 |
| Q5ZPR3 | CD276 | P30508 | HLA-C | Q06830 | PRDX1 | Q12931 | TRAP1 |
| P16070 | CD44 | P07910 | HNRNPC | P32119 | PRDX2 | P02766 | TTR |
| P60033 | CD81 | P32754 | HPD | Q13162 | PRDX4 | Q9BQE3 | TUBA1C |
| P23528 | CFL1 | Q86YZ3 | HRNR | Q9P2B2 | PTGFRN | P68366 | TUBA4A |
| Q9Y281 | CFL2 | P07900 | HSP90AA1 | P61026 | RAB10 | P07437 | TUBB |
| P06732 | CKM | P08238 | HSP90AB1 | P62491 | RAB11A | P68371 | TUBB4B |
| O00299 | CLIC1 | P0DMV8 | HSPA1A | P51153 | RAB13 | Q9BUF5 | TUBB6 |
| P09543 | CNP | P11021 | HSPA5 | P59190 | RAB15 | P0CG48 | UBC |
| P02452 | COL1A1 | P11142 | HSPA8 | Q15286 | RAB35 | Q9NQH7 | XPNPEP3 |
| P00450 | CP | P04792 | HSPB1 | P20339 | RAB5A | P07947 | YES1 |
| O75131 | CPNE3 | P05362 | ICAM1 | P51148 | RAB5C | P31946 | YWHAB |
| P01040 | CSTA | P17936 | IGFBP3 | P51149 | RAB7A | P62258 | YWHAE |
| P07339 | CTSD | P06756 | ITGAV | P63000 | RAC1 | P61981 | YWHAG |
| Q14118 | DAG1 | P05556 | ITGB1 | P11233 | RALA | Q04917 | YWHAH |
| P81605 | DCD | P05106 | ITGB3 | P11234 | RALB | P27348 | YWHAQ |
| | | | | | | P63104 | YWHAZ |

**Unique EMCV-$L^{Zn}$ (n=48)**

| Accession | Gene |
|---|---|
| P01023 | A2M |
| P02647 | APOA1 |
| P02652 | APOA2 |
| P06576 | ATP5F1B |
| P27824 | CANX |
| P40121 | CAPG |
| P47756 | CAPZB |
| Q00610 | CLTC |
| P68400 | CSNK2A1 |
| P00734 | F2 |
| P02774 | GC |
| P06396 | GSN |
| P09211 | GSTP1 |
| P04908 | HIST1H2AB |
| Q96KK5 | HIST1H2AH |
| P06899 | HIST1H2BJ |
| O60814 | HIST1H2BK |
| P68431 | HIST1H3A |
| P09651 | HNRNPA1 |
| P00738 | HP |
| P34932 | HSPA4 |
| P10809 | HSPD1 |
| Q06033 | ITIH3 |
| P01591 | JCHAIN |
| Q9C099 | LRRCC1 |
| P40926 | MDH2 |
| P35579 | MYH9 |
| O00159 | MYO1C |
| Q15365 | PCBP1 |
| Q86SE5 | RALYL |
| P27635 | RPL10 |
| P61313 | RPL15 |
| P18621 | RPL17 |
| Q07020 | RPL18 |
| P83731 | RPL24 |
| P36578 | RPL4 |
| P62917 | RPL8 |
| P62280 | RPS11 |
| P62851 | RPS25 |
| P62753 | RPS6 |
| P05109 | S100A8 |
| Q8NC51 | SERBP1 |
| P01009 | SERPINA1 |
| P29508 | SERPINB3 |
| P61956 | SUMO2 |
| P02787 | TF |
| Q86YD3 | TMEM25 |
| P18206 | VCL |

**Unique mock (n=39)**

| Accession | Gene |
|---|---|
| P23526 | AHCY |
| P05091 | ALDH2 |
| P78371 | CCT2 |
| P40227 | CCT6A |
| P13987 | CD59 |
| P14209 | CD99 |
| P60953 | CDC42 |
| Q14019 | COTL1 |
| P14384 | CPM |
| Q9H1C7 | CYSTM1 |
| Q14126 | DSG2 |
| O43854 | EDIL3 |
| P26641 | EEF1G |
| O43491 | EPB41L2 |
| Q96TA1 | FAM129B |
| P14324 | FDPS |
| O75223 | GGCT |
| Q8NBJ4 | GOLM1 |
| Q969P0 | IGSF8 |
| P17301 | ITGA2 |
| Q14847 | LASP1 |
| Q9NUP9 | LIN7C |
| Q08431 | MFGE8 |
| O95297 | MPZL1 |
| Q9Y623 | MYH4 |
| Q03181 | PPARD |
| P30041 | PRDX6 |
| P61019 | RAB2A |
| P61006 | RAB8A |
| Q92930 | RAB8B |
| Q15382 | RHEB |
| P62745 | RHOB |
| P84095 | RHOG |
| P06702 | S100A9 |
| Q99808 | SLC29A1 |
| Q8WUX1 | SLC38A5 |
| Q9P289 | STK26 |
| O43760 | SYNGR2 |
| Q9UN37 | VPS4A |

**Unique EMCV-Wt (n=21)**

| Accession | Gene |
|---|---|
| P15291 | B4GALT1 |
| Q5T0U0 | CCDC122 |
| O43633 | CHMP2A |
| Q9H444 | CHMP4B |
| Q9UBR2 | CTSZ |
| Q96JB1 | DNAH8 |
| Q969X5 | ERGIC1 |
| Q7Z5G4 | GOLGA7 |
| P47929 | LGALS7 |
| Q99732 | LITAF |
| Q12907 | LMAN2 |
| P20585 | MSH3 |
| Q15181 | PPA1 |
| P61106 | RAB14 |
| Q15293 | RCN1 |
| P10301 | RRAS |
| O00560 | SDCBP |
| P30626 | SRI |
| Q92734 | TFG |
| P22735 | TGM1 |
| Q9BVK6 | TMED9 |
| P12296 | EMCV-VP1 |
| P12296 | EMCV-VP2 |
| P12296 | EMCV-VP3 |
| P12296 | EMCV-2A |
| P12296 | EMCV-2C |
| P12296 | EMCV-3A |
| P12296 | EMCV-3C |

| Shared mock & EMCV-$L^{Zn}$ (n=37) | | Shared mock & EMCV-Wt (n=23) | | Shared EMCV-Wt & -$L^{Zn}$ (n=9) | |
|---|---|---|---|---|---|
| **Accession** | **Gene** | **Accession** | **Gene** | **Accession** | **Gene** |
| Q9UNQ0 | ABCG2 | P09525 | ANXA4 | P02649 | APOE |
| Q7Z5M8 | ABHD12B | P08133 | ANXA6 | P02749 | APOH |
| O60488 | ACSL4 | P20073 | ANXA7 | P63096 | GNAI1 |
| P23634 | ATP2B4 | P08174 | CD55 | P14625 | HSP90B1 |
| P61769 | B2M | P19256 | CD58 | P19823 | ITIH2 |
| P01024 | C3 | P21926 | CD9 | P35268 | RPL22 |
| P52907 | CAPZA1 | Q86YQ8 | CPNE8 | P18077 | RPL35A |
| P12277 | CKB | P78310 | CXADR | P62263 | RPS14 |
| P00533 | EGFR | P31025 | LCN1 | Q5T750 | XP32 |
| P60842 | EIF4A1 | P07948 | LYN | | |
| P13929 | ENO3 | P23284 | PPIB | | |
| P62805 | HIST1H4A | P15151 | PVR | | |
| Q00839 | HNRNPU | P10114 | RAP2A | | |
| P01112 | HRAS | O75695 | RP2 | | |
| P46940 | IQGAP1 | P05388 | RPLP0 | | |
| P17931 | LGALS3 | O14828 | SCAMP3 | | |
| Q92542 | NCSTN | O75396 | SEC22B | | |
| P06748 | NPM1 | Q96P63 | SERPINB12 | | |
| P09874 | PARP1 | P50454 | SERPINH1 | | |
| P30086 | PEBP1 | O00161 | SNAP23 | | |
| P52209 | PGD | Q12846 | STX4 | | |
| O15031 | PLXNB2 | P49755 | TMED10 | | |
| P62937 | PPIA | P10599 | TXN | | |
| Q9H0U4 | RAB1B | | | | |
| P50914 | RPL14 | | | | |
| P46776 | RPL27A | | | | |
| P49207 | RPL34 | | | | |
| P62249 | RPS16 | | | | |
| P62854 | RPS26 | | | | |
| P61247 | RPS3A | | | | |
| P62241 | RPS8 | | | | |
| Q9H2H9 | SLC38A1 | | | | |
| O14745 | SLC9A3R1 | | | | |
| Q8IZP2 | ST13P4 | | | | |
| P37802 | TAGLN2 | | | | |
| P06753 | TPM3 | | | | |
| P04004 | VTN | | | | |
